## Supplementary Materials for "Information about task duration influences energetic cost during split-belt adaptation and retention of walking patterns post-adaptation"

### Information about task duration influenced heart rate, and RPE during split-belt adaptation

The average heart rate was 114 ± 17 bpm during EA, 109 ± 18 during MA, and 113 ± 18 during LA, indicating that the split-belt adaptation task was of moderate cardiovascular intensity. The linear model assessing heart rate as a function of group, time, and baseline heart rate during split-belt adaptation returned a significant intercept (β_0_=106, 95%CI [100, 112], p<0.001), a significant main effect of baseline heart rate (β_Base_=12, 95%CI [7, 16], p<0.001) and a significant main effect of True Group (β_GroupT_=11, 95%CI [3, 19], p=0.006) and of False Group (β_GroupF_=9, 95%CI [1, 18], p=0.024) indicating higher heart rates in the True and False groups relative to the Control Group (adjusted R^2^=0.66) (Fig. 3D).

The linear model assessing RPE as a function of group, time, and baseline RPE during split-belt adaptation returned a significant intercept (β_0_=3.4, 95%CI [3, 4], p<0.001), a significant main effect of baseline RPE (β_Base_=0.59, 95%CI [0.15, 1], p=0.009) and a significant interaction between the True group and baseline RPE (β_GroupT:BaseRPE_=0.70, 95%CI [0.23, 1.16], p=0.003) (Fig. 3E). Estimated marginal trends of the slope between base RPE by groups show a greater slope in the True group compared to both the Control group (slope True = 1.32 vs slope Control = 0.54, p=0.005) and the False group (slope False = 0.63, p=0.004), indicating a greater increase in RPE relative to baseline in the individuals in the True group who have detailed information about time remaining (adjusted R^2^=0.39).

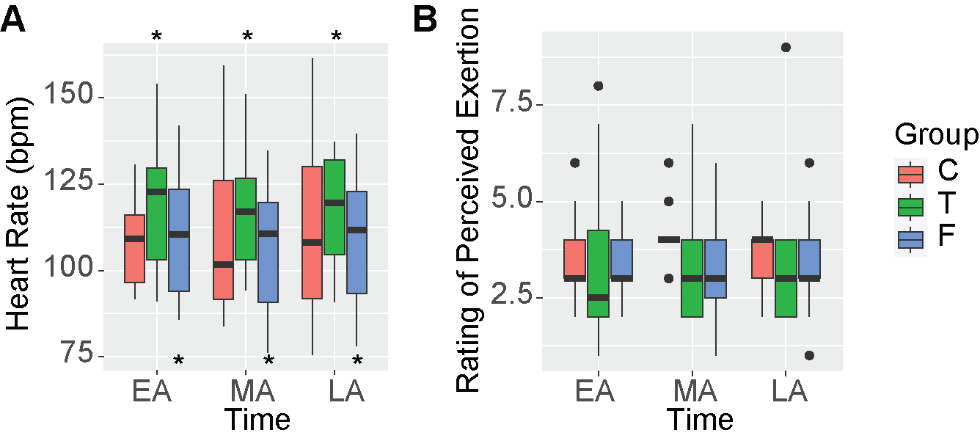

**Supplementary Figure 1.** A) Average heart rate during EA, MA, and LA. A significantly higher heart rate was seen in the True group relative to Control (p=0.006) and in the False group relative to Control (p=0.024). B) Rating of perceived exertion as indicated by participants during EA, MA, and LA. A greater increase in RPE from baseline was observed in the True group.

#### **Information about task duration influenced negative work during split-belt adaptation**

The linear model assessing negative work by the fast leg as a function of group, time and baseline work during split-belt adaptation returned a significant intercept (β_0_=-.010, 95%CI [-0.014, -0.007], p<0.001), a significant interaction between time and baseline negative work indicating that negative work changed proportionally to baseline during adaptation (β_TimeMA:Base_=0.0032, 95%CI [0.00, 0.006], p=0.023 and β_TimeLA:Base_=0.0038, 95%CI [0.001, 0.007], p=0.006). We also observed a significant interaction in the True group (β_GroupT:Base_=0.0033, 95%CI [0.00, 0.006], p=0.021) indicating that the True group generated less negative work (adjusted R^2^=0.18) (Fig. 4A, E). Thus, we observed that individuals in the True group generated less negative work with the fast leg, opposite to the strategy shown to allow taking advantage of work from the treadmill to reduce work by the legs and reduce metabolic cost [1,2].

The linear model assessing negative work by the slow leg as a function of group, time and baseline work during split-belt adaptation returned a significant intercept (β_0_=-0.020, 95%CI [-0.022, -0.016], p<0.001), a significant main effect of baseline work by the slow leg (β_Base_=0.0037, 95%CI [0.001, 0.0006], p=0.002) and a significant effect of time (β_TImeMA_=0.0097, 95%CI [0.005, 0.014], p<0.001 and β_TimeLA_=0.011, 95%CI [0.006, 0.016], p<0.001) (adjusted R^2^=0.44). This model indicates that across all groups, all participants decreased negative work by the slow leg during adaptation (Supplementary Fig. 2).

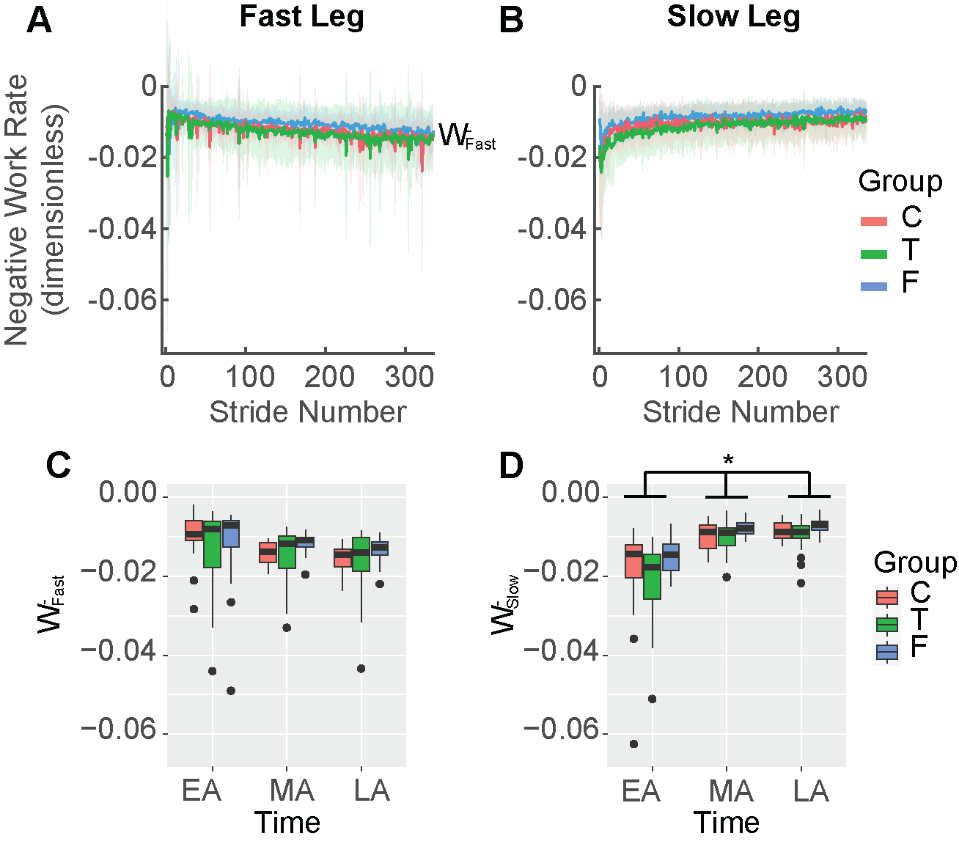

**Supplementary Figure 2. Negative work by the fast and slow legs during split-belt adaptation.** A) Timeseries of negative work by the fast leg. B) Timeseries of negative work by the slow leg. C) Negative work by the fast leg during the 5 strides corresponding to EA, MA, and LA. D) Negative work by the slow leg between groups during the 5 strides corresponding to EA, MA, and LA. We observed a significant reduction in work from EA to MA and LA across groups. W^-^: negative work.

#### **Information about task duration during split-belt adaptation influenced metabolic cost during post-adaptation**

The linear model assessing metabolic cost as a function of group, time, and baseline cost during post-adaptation returned a significant intercept (β_0_=2.59, 95%CI [2.39, 2.79], p<0.001), a significant main effect of baseline metabolic cost (β_Base_=0.21, 95%CI [0.06, 0.37], p=0.006), a significant main effect of LP (β_TimeLP_=-0.34, 95%CI [-0.62, -0.06], p=0.016), and a significant interaction in the True group (β_GroupT:BaseMet_=0.34, 95%CI [0.15, 0.53], p<0.001). Estimated marginal means of the slope by groups show a greater slope in the True group (slope T = 1.1) compared to both The Control group (slope C = 0.38, p=0.002) and The False group (slope F = 0.59, p=0.015), indicating a greater increase from baseline metabolic cost during the post-adaptation task in the True group (adjusted R^2^=0.52) (Supplementary Figure 3).

No differences in heart rate were observed between groups or over time during post-adaptation. The RPE did not change over time (β_0_=2.25, 95%CI [1.8, 2.7], p<0.001), but the True group showed a significant interaction with baseline (β_GroupT:BaseRPE_=0.70, 95%CI [0.07, 1.03], p=0.003). Estimated marginal trends of the slope between base RPE and RPE by groups did not show significant differences between groups (adjusted R^2^=0.39).

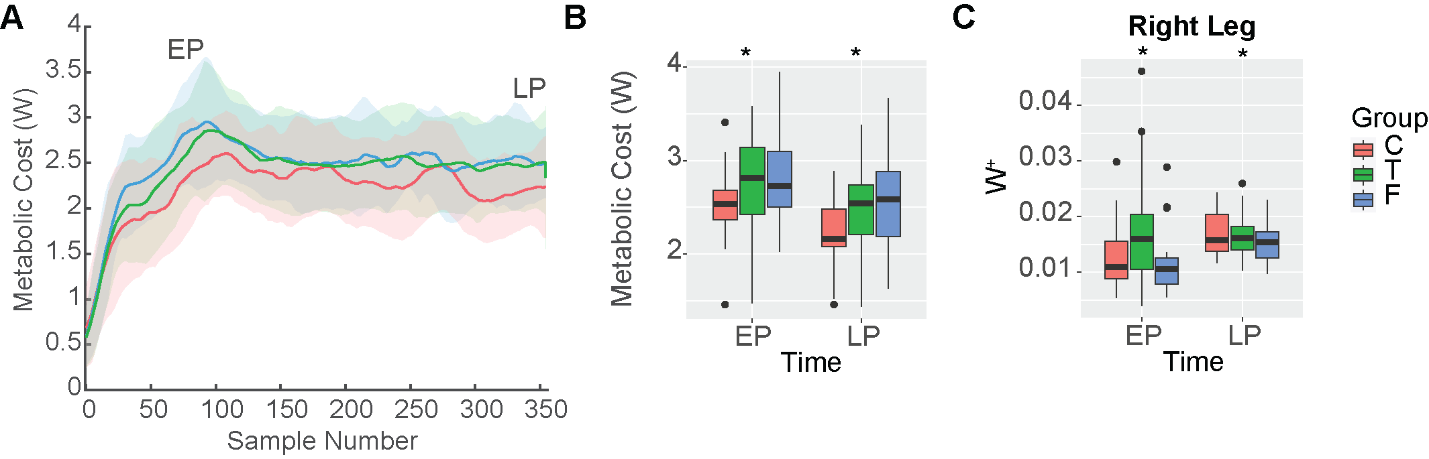

**Supplementary Figure 3. Metabolic cost and work during post-adaptation.** A) Net metabolic power timeseries during post-adaptation between the control (C – red), true knowledge of task duration (T – green) and false knowledge of task duration (F - blue) groups. Time windows indicating EP, and LP are indicated in the figure. B) Average metabolic cost during EP, and LP. A higher cost was observed in the True group (p<0.050). C) Average positive work by the right leg during EP and LP. Participants in the True group maintained the higher work with the right leg (p=0.009).

#### **Information about task duration of split-belt adaptation influenced work during post-adaptation**

The linear model assessing positive work by the right leg (previously on the slow belt) as a function of group, time, and baseline work during post-adaptation returned a main effect of baseline work by the right leg (β_Base_=1.01, 95%CI [0.73, 1.29], p<0.001), and a significant main effect of The True group (β_GroupT_=0.005, 95%CI [0.0012, 0.008], p=0.009). These results indicate that in the True group, positive work was higher during post-adaptation than in the C and F groups (adjusted R^2^=0.38) (Supplementary Figure 3C). No differences between groups were observed for positive work by the left leg and negative work by the left and right legs.
